# Degenerating *Drosophila* Larval Epidermal Cells Drive Thorax Closure

**DOI:** 10.1101/2020.09.22.304451

**Authors:** Thamarailingam Athilingam, Saurabh Singh Parihar, Rachita Bhattacharya, Mohd. Suhail Rizvi, Amit Kumar, Pradip Sinha

**Affiliations:** Department of Biological Sciences and Bioengineering, Indian Institute of Technology Kanpur, India, 208016; Mechanobiology Institute, National University of Singapore, 5A Engineering Drive 1, Singapore 117411; Indian Institute of Technology Hyderabad, India 502285; Albert Einstein College of Medicine, 1300 Morris Park Avenue, Bronx, New York

**Keywords:** Thorax closure, Larval epidermis, Organogenesis, *Drosophila*

## Abstract

Adult thorax formation in *Drosophila* begins during pre-pupal development by fusion of its two contralateral progenitor halves, the heminotal epithelia (HE). HEs migrate and replace an underlying cell layer of thoracic larval epidermal cells (LECs) during a morphogenetic process called thorax closure. The LEC layer has so far been proposed to be a passive substrate over which HEs migrate before their zipping. By contrast, here we show that the pull forces generated within the LEC layer drive HE migration. During thorax closure, the LECs display actomyosin-mediated contraction, via enrichment of non-muscle myosin-II and actin, besides squamous-to-pseudostratified columnar epithelial transition and tissue shrinkage. This shrinkage of the LEC layer is further accompanied by cell extrusion and death, that prevent overcrowding of LECs, thereby promoting further shrinkage. The pull forces thus generated by the shrinking LEC layer are then relayed to the HEs by their mutual adhesions via βPS1 (Mys) and αPS3 (Scb) integrins. Suppression of cell death in the LEC layer by a gain of p35 leads to cell overcrowding, which impedes HE migration and zipping. Further, knockdown of *sqh*, the light chain of non-muscle myosin II, in LECs or integrins (*mys* or *scb*) in either the LEC layer or in the HEs, or both abrogate thorax closure. Mathematical modeling also reveals the biophysical underpinnings of the forces that drive this tissue closure process wherein a degenerating LEC layer mediates its succession by the future adult primodia. These essential principles of thorax closure appear ancient in origin and recur in multiple morphogenetic contexts and tissue repair.

## Introduction

During larval-pupal molt in holometabolous insects, a high ecdysone hormone titer triggers the replacement of larval structures by their adult primordia (for a recent review, see (Truman and Riddiford, 2019)). Thorax closure in *Drosophila* exemplifies one such morphogenetic event where larval epidermal cells (LECs) of the three thoracic body segments are replaced by their corresponding adult primordia. Adult thorax (or notum) is formed from three paired imaginal progenitors: namely, dorsal prothoracic (humeral) discs, proximal cells of the mesothoracic wing discs, and the metathoracic haltere imaginal discs (for review, see (Beira and Paro, 2016; Fristrom and Fristrom, 1993)). These bilateral precursors of the adult thorax, following their eversion (Pastor-Pareja et al., 2004), undergo fusion with each other to form two contralateral heminotal epithelia (HE), one each on the left and the right thoracic region of the prepupa. Thorax closure ensues (Fristrom and Fristrom, 1993) when these two opposing contralateral HEs migrate, replacing the LEC layer and zip at the future midline of the adult body: a culmination of an elegant process of epithelial tissue closure (see Figure S1A) (Agnès et al., 1999; Kumar et al., 2020; Martín-Blanco et al., 2000; Usui and Simpson, 2000; Zeitlinger and Bohmann, 1999).

Historically, studies on thorax closure (Agnès et al., 1999; Martín-Blanco et al., 2000; Usui and Simpson, 2000; Zeitlinger and Bohmann, 1999) closely followed the leads obtained from another epithelial tissue closure event in *Drosophila*: namely, embryonic dorsal closure (DC) (for review, see (Belacortu and Paricio, 2011)). Like in thorax closure, during DC too, two lateral embryonic epidermal cell sheets converge dorsally, replacing an extra-embryonic flattened epithelial cell layer, the amnioserosa (for recent reviews, see (Hayes and Solon, 2017; Kiehart et al., 2017)). Anatomical parallels could be drawn between the embryonic amnioserosa and the thoracic LEC layer as both degenerate and are replaced during their respective tissue closure events. Likewise, lateral embryonic epidermis and pre-pupal HEs represent the cell layers that migrate during DC and thorax closure, respectively. The leading edges (LEs) of the lateral embryonic epidermis have been proposed to mediate DC based on their supracellular actin cables via their purse-string-like contractions (Kiehart et al., 2000), while actin-rich filopodial projections from the LEs contribute to their zipping (Jacinto et al., 2002). Given these precedences in DC, previous studies in thorax closure also explored the LEs of HEs as potent hubs of mechano-regulation and cell signaling (for review see, (Belacortu and Paricio, 2011; Harden, 2002)). Indeed, it was seen that perturbations in signaling pathways such as JNK, D-Fos, or Dpp in the LEs, affect thorax closure, reminiscent of similar outcomes in DC (for review, see (Belacortu and Paricio, 2011; Harden, 2002; Martín-Blanco and Knust, 2001)). Further, LEs of HEs also displayed actin-rich filopodia (Agnès et al., 1999; Martín-Blanco et al., 2000; Pastor-Pareja et al., 2004; Usui and Simpson, 2000; Zeitlinger and Bohmann, 1999), that are known to display reversible contacts with substratum in various morphogenetic contexts (Parsons et al., 2010). It was therefore proposed that contralateral HEs crawl over the LEC layer, wherein the underlying LECs are left ‘below and behind and eventually delaminate from the edges’(Martín-Blanco et al., 2000). In essence, these interpretations of thorax closure present the thoracic LEC layer as a scaffold over which HEs migrate (see Figure 6 in (Martín-Blanco et al., 2000)). Indeed, somewhat comparable roles LEs play in other organisms, such as gastrulation in sea urchin embryo (Gustafson, 1963), ventral enclosure in *C. elegans* (Williams-Masson et al., 1997), or closure of epithelial gaps during wound healing (Martin, 1997; Martin and Lewis, 1992) presented a framework wherein LEs of migrating HEs appeared as the drivers of thorax closure.

Subsequent studies on DC, however, led to a substantial revision in the understanding of its mechanism wherein amnioserosa—rather than the LEs of the lateral embryonic epidermis—was shown to be the principal source of forces for tissue closure. For instance, amnioserosa shrinks by apoptosis (Toyama et al., 2008) and displays non-muscle myosin II-dependent pulsed contractions (Franke et al., 2005; Solon et al., 2009), which collectively generate forces that drive migration of the lateral embryonic epidermis. Strikingly, more recent studies have shown that pull forces generated by the amnioserosa layer alone are sufficient to drive DC (Ducuing and Vincent, 2016; Pasakarnis et al., 2016), while supracellular actomyosin cables in the LEs of the lateral embryonic epidermis display ancillary roles in dorsal closure: such as maintenance of epithelial cell packing, tissue organization, and polarity (Ducuing and Vincent, 2016) besides aiding in scar-free zipping (Pasakarnis et al., 2016). In essence, these findings assign degenerating amnioserosa as the major force-generating tissue during DC (for review, see (Hayes and Solon, 2017; Kiehart et al., 2017)).

In the light of these emergent understandings of DC, here we have re-examined the roles of a degenerating LEC layer in thorax closure. We show that non-muscle myosin II-mediated contractions along with cell extrusions and cell death in the LEC layer, generate the pull forces that are then relayed to the flanking HEs due to their integrin-based adhesions. Our mathematical modeling further formalizes these biophysical principles underlying LECs-driven tissue closure event, wherein a degenerating, yet dynamic LEC layer drives its succession by the future adult primordia.

## Results

### Progressive shrinkage of the LEC layer during thorax closure

In *Drosophila*, thorax closure sets in at an early stage of pupal development—often referred to as ‘pre-pupal’ stage (0-12 hrs <u>a</u>fter <u>p</u>uparium formation (APF) (Fristrom and Fristrom, 1993))—that lasts from 4.30 to 6.00 hrs APF at 25□C (Kumar et al., 2020; Martín-Blanco et al., 2000; Usui and Simpson, 2000; Zeitlinger and Bohmann, 1999). Leading-edge (LE) cells of the contralateral HEs—which are seen in close apposition with the underlying LEC layer (Martín-Blanco et al., 2000; Pastor-Pareja et al., 2004; Usui and Simpson, 2000)—display upregulated expression of a JNK reporter, *puc-lacZ* (Figure 1A, A’), besides actin-enriched filopodial and lamellipodial protrusions (red and blue arrows, respectively, Figure 1A’).

**Figure 1.**
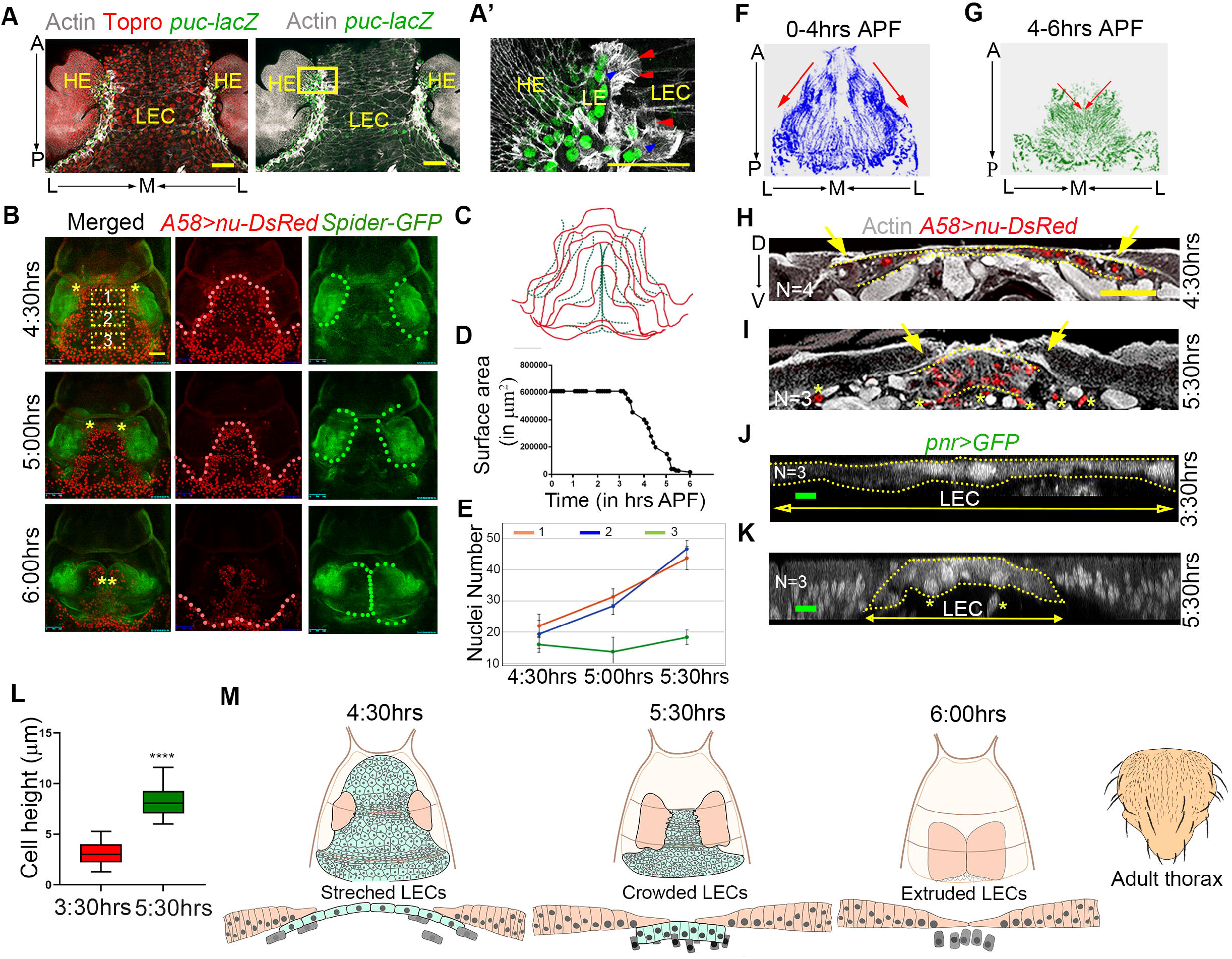
LEC layer dynamics during thorax closure. **(A-A’)** HE and LEC layer during the course of thorax closure: *puc-lacZ* (β-gal, green) mark the leading edge; F-actin (grey), and nuclei (red) (A). (A’) Magnified box of (A) reveals lamellipodia (blue arrowhead) and filopodia (red arrowhead). Scale bar: 50 μm. **(B-C)** Snapshots (from left panel of Video 1) for the progression of thorax closure; the edges of HE (green outlined) and LEC layer (red outline) are also shown separately in (C). Scale bar: 100 µm. **(D)** Change in surface area of LEC layer before and during the course of thorax closure (quantified from left panel in Video 1) (N=1). **(E)** Increase in the number of nuclei—to predict overcrowding—in the medial region of LEC layer in T2 (Box 1 and 2) segment as compared to T3 segment (Box 3) (see Figure1B for boxes location) (N =2; Mean ± SD). **(F-G)** Tracks of cell movement—as traced from their nuclei (from right panel of Video 1)—in the LEC layer before (0-4 hrs, blue, F) and during (4-6 hrs, green, G) thorax closure. **(H-I)** Cryo-cross sections of T2 segment at 4:30 hrs (H) and 5:30 hrs (I) APF, stained for actin (grey) and LECs (red nucleus) (yellow dotted outlined), the HE (yellow arrows) and delaminating nuclei (yellow star) (N =number or samples tested), Scale bar: 50 µm. **(J-L)** Cell height of3:30 hrs (early) (J) and 5:30 hrs (late) (K) of LEC layer (yellow dotted outline), with delaminated cells (yellow stars); and their respective quantification (L) (N=3 for each of the stages with 10 independent measurements from each sample, Student’s t-test, ****p<0.0001, Mean± SEM). Scale bar: 5 μm. (M) Cartoon representation of the dynamic changes in the LEC layer (blue-green)—from a stretched to a crowded state followed by the cell extrusion—parallel to HE migration (brown) during thorax closure.

We first re-examined the LEC layer and HE movements and dynamics during thorax closure by live-cell imaging. Nuclei of LECs were selectively marked (*A58*>*nu-DsRed*, also see Figure S1D) (Anderson and Galko, 2014; Galko and Krasnow, 2004) along with Spider-GFP—reporter for *gish* gene (Morin et al., 2001)—which marks the cell boundaries of both HEs and LECs (Morin et al., 2001) (left panel in Video 1, Figure 1B). We noticed a progressive shrinkage of LEC layer perimeter (left and right panel in Video 1, Figure 1B, and red outlines in Figure 1C)— coincident with the progression of migrating LEs of the contralateral HEs (left panel in Video 1, Figure 1B and green outlines in Figure 1C); with LEC layer displaying a comparable rate of shrinkage (4.1±0.77 µm/min, Figure 1C) to that of HEs migration (4.3±0.46 µm/min for HE, Figure 1C). This shrinkage in the LEC layer results in a precipitous drop in its surface area during thorax closure (4:30 to 6:00 hrs APF, Figure 1D) besides nearly a two-fold increase in its cell density, as seen from nuclear counts in its medial region (Figure 1E, see representative boxes 1-3 in Figure 1B).

Trajectories of LECs migration—visualized by tracking their nuclear displacements (right panel in Video 1) revealed spatio-temporal characteristics: initial trajectories are aligned along the anterior-to-posterior (A→P) body axis (between 0-4 hrs APF; right panel in Video 1, Figure 1F): that is, when thoracic LECs display apolysis or detachment from the overlaying pupal cuticle (Usui and Simpson, 2000) while later, during thorax closure proper, these trajectories were seen along the L→M axis (4:30-6:00 hrs APF; right panel in Video 1, Figure 1G), parallel to the direction of HEs migration (left panel in Video 1; Figure 1B).

Cryo-cross sections along the second thoracic pupal segment (T2) further revealed that LEs of HEs remain in contact with the lateral edges of the LEC layer (Figure 1H-I) (also see (Usui and Simpson, 2000)). Further, we noticed a transition of the LEC layer from flattened squamous cells, with widely-spaced nuclei aligned in a single plane (4:30 hrs APF, red nuclei in Figure 1H) to densely packed tall columnar cells wherein nuclei are seen at different planes along their dorso-ventral axis (5.30 hrs APF, red nuclei in Figure 1I), revealing their transition to a pseudo-stratified state (also see (Norden, 2017)). Moreover, at later stages, LECs were also seen to be extruded from the layer (yellow stars in Figure 1I). Confocal microscopic optical X-Z sections of the LEC layer further confirmed an increase in its cell packaging density (Figure 1K compared to 1J), increased cell heights (Figure 1K, L) with accompanying pseudo-stratification (Figure 1K), and, finally, cell extrusion (yellow stars, Figure 1K) during thorax closure.

Together, these observations reveal a host of novel features of epithelial dynamics in the LEC layer, which present a compelling argument for a possible link for its active role in thorax closure (Figure 1M).

### Delamination and cell death in the LEC layer facilitate unimpeded HE migration and zipping

Programmed cell death (PCD) is a key event in early embryonic development in both invertebrates and vertebrates (for reviews, see (Nagata, 2018; Singh et al., 2019; Suzanne and Steller, 2013; Teng and Toyama, 2011)). For instance, during DC, PCD in the amnioserosa contributes to generating pull forces that drive tissue closure (Teng and Toyama, 2011; Toyama et al., 2008). Likewise, elimination of abdominal LECs by apoptosis prefigures the expansion of histoblast nests (Ninov et al., 2007; Teng and Toyama, 2011) via a ‘push-pull coordination’ with the latter (Prat-Rojo et al., 2020). It is therefore conceivable that delamination in LECs (Figure 1I, K) and their eventual loss by apoptosis (Ninov et al., 2007) could also be crucial for thorax closure by removing the impediment of cell crowding on the path of migrating HEs.

A careful examination of Spider-GFP marked LECs (Figure S2A) revealed their shrinking perimeters (yellow arrows, Figure S2A’) while their live imaging using Lachesin-GFP—a septate junction protein reporter (Morin et al., 2001)—further revealed progressive apical constrictions (yellow and red arrows in Video 2 and Figure 2A) and delamination (Figure S2B). Previously, it has been shown that delaminating cells often display apoptosis, marked by caspase activation (Muliyil et al., 2011; Saias et al., 2015; Teng et al., 2017). Indeed, an Apoliner construct—a genetically encoded caspase sensor and an early apoptotic marker (Bardet et al., 2008)—revealed early caspase activation in the majority of LECs (Figure S2C), which reconciles well with the fact that most LECs are delaminated (Figure S2D), leading to their elimination by cell death.

**Figure 2:**
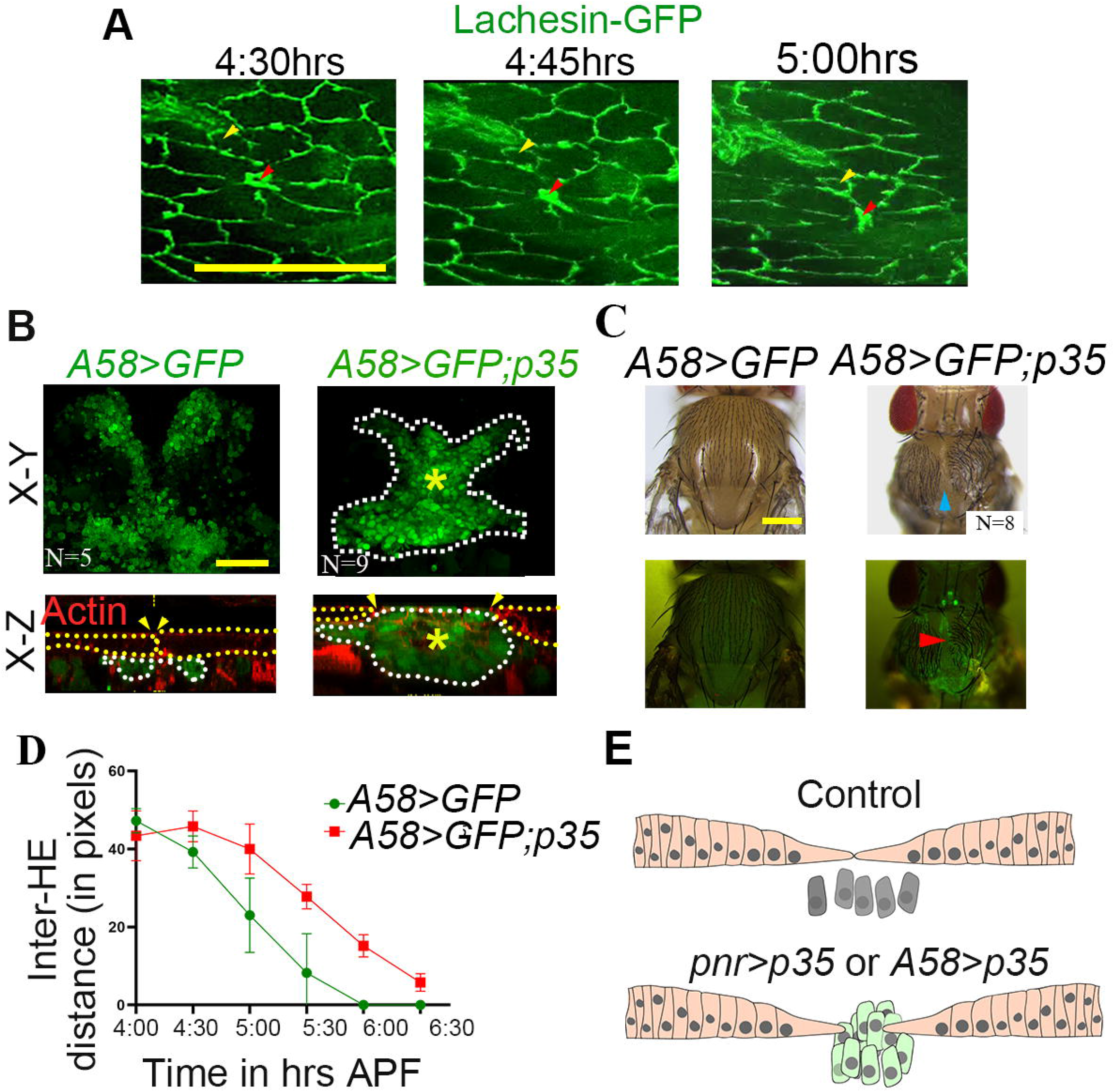
Cell delamination and death in LEC layer during thorax closure. **(A)** Snapshots of Lachesin-GFP marked cell outline of LECs (also see Video 2) showing cells undergoing apical constriction (yellow arrowhead) and delamination (red arrowhead) (Scale bar: 25 μm). **(B)** Top and X-Z view for controls compared to *p35* overexpression using *A58-Gal4* (also see Figure S1D). Outlines of HE (yellow dotted lines) and LEC layer (white dotted lines and counterstained for actin (red, shown only in lower panel) to mark cell cytoskeleton and muscles. Large masses of LECs shown as (yellow stars) (Scale bar: 50 μm). **(C)** Adult thorax (upper panels) and their respective fluorescent images (lower panels) from the above case (B). Cleft thoraces phenotypes following *p35* overexpression (blue arrowheads in upper panel) and red arrowheads mark persistence of GFP-marked LECs in adults (lower panel) (Scale bar: 200 μm). **(D)** Inter-HE distances in case of *A58>GFP* (control, green) and their respective *p35* overexpression (*A58>GFP; p35*, red) (N=3 for each case, Mean ± SEM, two-way ANOVA was performed, **** p value: <0.0001). **(E)** Cartoon representation of cross-sectional view of the control and those displaying *p35* over-expressions at 6 hrs APF.

To further probe the role of cell death in the LEC layer, we examined the consequences of gain of p35—an inhibitor of caspase-3-like proteins in *Drosophila* (Hay et al., 1994)—by selectively driving *UAS-p35* transgene under the *A58-Gal4* driver (Figure S1D). Comparison of the endpoint of thorax closure, as revealed by X-Y (upper panel, Figure 2B) and X-Z optical sections (lower panel, Figure 2B), at 6:00 hrs APF for the control (*A58>GFP*, Figure 2B) and in case of *p35* overexpression (*A58>GFP; p35*, Figure 2B) revealed cell overcrowding (yellow star, Figure 2B, lower panel) in the latter with an accompanying impediment in HE fusion, as seen from their failed zipping and persistence of large inter-HE gap (arrowheads in Figure 2B, lower panel). Not surprisingly, adults emerging out of these pupae displayed a characteristic cleft-thorax phenotype (arrowhead, Figure 2C) besides the persistence of non-extruded LECs in the freshly eclosed adults (arrowhead, GFP, Figure 2C) and prolongation of the duration of thorax closure (Figure 2D). Similar overcrowding and failure in HE fusion can also be seen in the case of *p35* overexpression both in HEs and LECs (Figure S3A) using the *pnr-Gal4* driver (Figure S1B). These adults also displayed the characteristic cleft-thorax phenotype (arrowhead in the upper panel of Figure S3B) and persistence of non-extruded LECs in freshly eclosed adults (arrowhead in the lower panel of Figure S3B), besides prolonging the duration of thorax closure (Figure S3C). By contrast, a selective overexpression of *p35* in only the HEs, using *332*.*3Gal80; pnr-Gal4* (see Figure S2D) did not affect thorax closure (Figure S3D).

Together, these results reveal that cell death in the LEC layer aid in its shrinkage, help reduce overcrowding of LECs and, facilitate HE migration and fusion during thorax closure (Figure 2E).

### Non-muscle myosin-II-mediated contraction in LEC layer drives migration of HEs

Given the possible active contraction (Video 2, Figure 2A) and overall shrinkage in the LEC layer (Video 1, Figure 1), it is likely that during thorax closure it can generate pull forces that facilitate HE migration. To test this possibility, we first ablated the LEC layer using a short pulse of the laser during thorax closure and visualized the fallouts (red arrow in Video 3). Indeed, ablation of the LEC layer during thorax closure brings HE migration to a halt, which then recedes away (red stars in Figure 3B compared to 3A, Video 3 compared to left panel in Video 4) as seen from a rapid increase in their inter-HE distance (red stars, Figure 3B and C). This fallout of the laser ablation in the LEC layer, therefore, suggests that the LEC layer is under tension and can generate pull forces. Conversely, it may also be argued that laser ablation disrupts the LEC layer, thereby disrupting the scaffold over which contralateral HE migrates (Martín-Blanco and Knust, 2001; Martín-Blanco et al., 2000).

**Figure 3.**
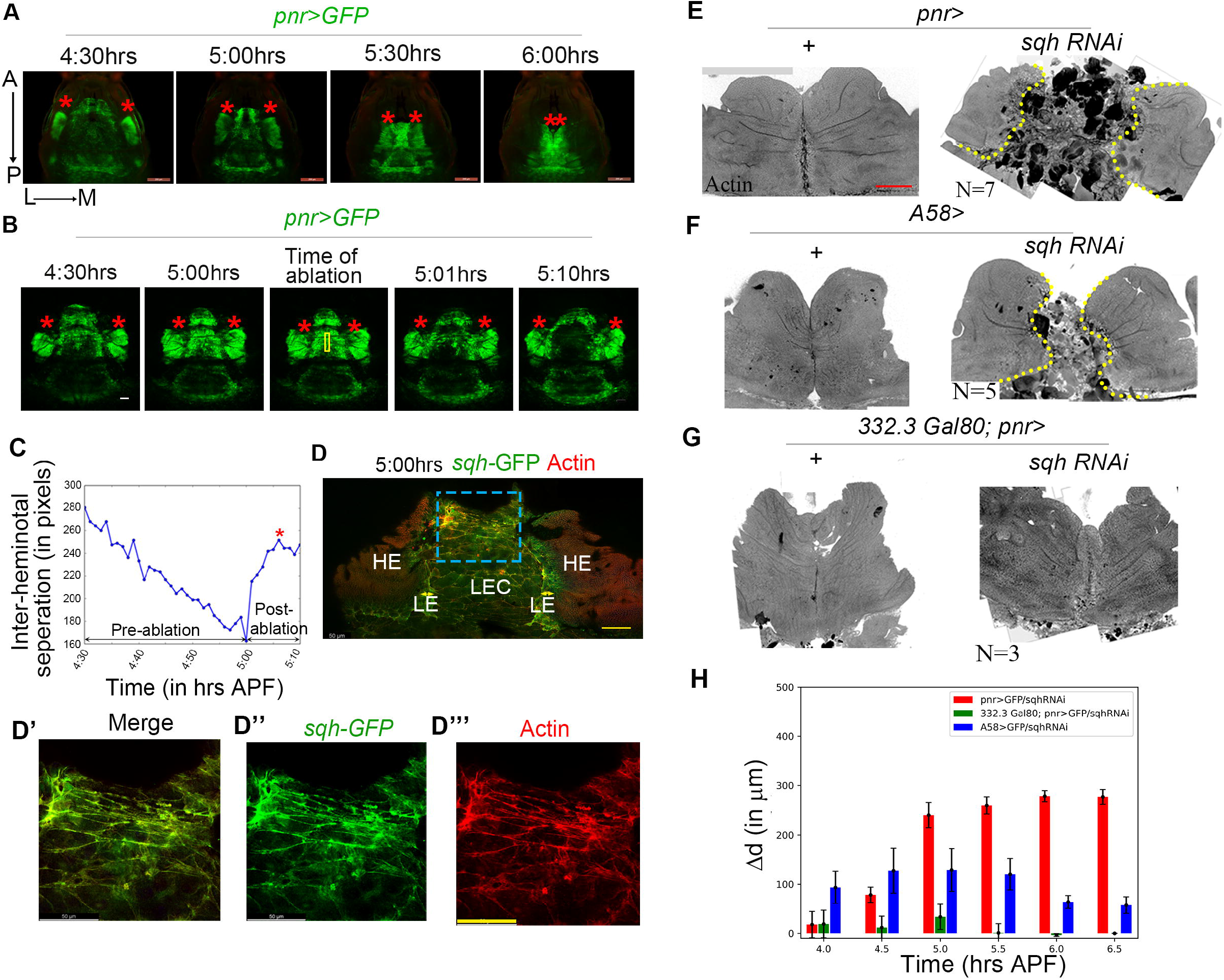
Non-muscle myosin II in LEC layer drives thorax closure. **(A-B)** Time-lapse snapshots of thorax closure in *pnr>GFP* (left panel, Video 4) (A) and following laser ablation (B) (Video 3). Yellow box in (B) represents the region of interest (ROI) for laser ablation. Red star marks the tips of the contralateral HE (Scale bar: A-200 µm and B-50 µm). **(C)** Graphical plot to show the increased inter-HE distances following laser ablation (B) during thorax closure. **(D-D’’’)** Sqh expression in the migrating HE and LEC layer, counterstained with actin (red). The blue boxed area of (D) is magnified (D’) to display the patterns of enrichments of Sqh*-*GFP (D’’) and actin (D’’’) in the LEC layer (Scale bar: 50 μm). **(E-G)** Whole-mounts preparation at 6 hrs APF for control (+, left panel) and their respective *sqh* knockdown (*sqh*-RNAi, right panel) using various Gal4s (see Figure S1B-D); yellow dotted lines represent the edges of HE that failed to zipper (Scale bar: 50 μm). **(H)** Deviation of inter-HE separations at specific time points between control and *sqh* knockdown in E-G (see Video 5 with their respective panels in Video 4) (N =4; Mean ± SEM, ANCOVA test was performed to compare the three linear regression models fits to the 3 cases; ***, p values: <0.001).

To distinguish between the two alternative roles of the degenerating thoracic LEC layer during thorax closure: namely, its role as a passive scaffold versus active contractile force generator, we further asked if, like the contractile amnioserosa cells (Franke et al., 2005; Young et al., 1993), LECs, too, display non-muscle myosin II-dependent contractions. We examined the expression of the regulatory light chain of non-muscle myosin II, *spaghetti squash (sqh)*—based on its GFP reporter, Sqh-GFP (Sisson et al., 2000)—during the thorax closure. Sqh regulates non-muscle myosin II contractile functions via its phosphorylation (Vereshchagina et al., 2004). We noticed an elevated Sqh-GFP level in LECs (Figure 3D-D’’) and also in the LEs of the contralateral HE (Figure 3D), reminiscent of the domains of expression of Zip/Myo-II in analogous cell types in the embryonic dorsal closure: that is, amnioserosa and the LE of the lateral epidermal layer, respectively (Franke et al., 2005). LECs also displayed enrichment of actin, which co-localized with Sqh*-*GFP on the cell edges and also in their cytoplasm (Figure 3D-D”’) (Franke et al., 2005; Gorfinkiel, 2016).

To assess the functional relevance of Sqh enrichment in the LEC layer during thorax closure, we knocked down *sqh* simultaneously in both the LEC layer and the HE (Figure S1B) (left panel in Video 5 compared to left panel in Video 4), selectively in only the LEC layer (Figure S1D) (middle panels in Video 5 compared to the middle panel in Video 4) or in the HE (Figure S2C) (right panel in Video 5 compared to the right panel in Video 4). We noticed a failure in thorax closure when the *sqh*-knockdown spanned the LEC layer (double arrow in left and middle panel in Video 5, Figure 3E and 3F). Quantification of the inter-HE distances at various time intervals of thorax closure following *sqh* knockdown in the LECs or both in LECs and HEs, further revealed persistent gaps and failed closure (red and blue bars in Figure 3H). We also note that knockdown of *sqh* in HEs does not have much impact on thorax closure (right panel in Video 5, Figure 3G and green bars in Figure 3H), reminiscent of the possible redundant role of LEs in DC ((Franke et al., 2005) for review see (Kiehart et al., 2017)).

Thus, during thorax closure, as in DC, pull forces generated by non-muscle myosin II from the LECs drive the HE migration, which is reminiscent of the role played by amnioserosa during dorsal closure (for review, see (Kiehart et al., 2017)).

### Pull forces from the LEC layer are relayed to the contralateral HE via their integrin-based adhesion

Adhesion between two apposed cell layers can impact their respective morphogenesis and mechano-transduction (Sun et al., 2016). Such inter-tissue adhesions are often mediated by the members of the integrin family, comprising of α and β subunits of position-specific (PS) integrins (Narasimha and Brown, 2013). For instance, during embryonic DC, the lateral embryonic epidermal layers adhere to the centrally placed amnioserosa via enrichment of βPS1 (Myospheroid, Mys) and αPS3 (Scab, Scb) dimers at their tissue interfaces, thereby facilitating relay of forces between these two cell layers (Narasimha and Brown, 2004).

We thus examined the fallouts of knockdowns of integrins during thorax closure. *Drosophila* genome encodes 2β (βPS1 and βυ) and 5α (αPS1-5) subunits of integrins that form multiple versions of heterodimeric (αβ) integrin complexes in different contexts of embryonic and adult development (Narasimha and Brown, 2004; Narasimha and Brown, 2013). Individual knockdowns using RNAi constructs of 2β and 4α (αPS1-4) integrins in both HE and LECs (Figure S1B) showed very high pupal lethality upon *mys (β-PS1)* or *scb (α-PS3)* knockdown, suggesting their possible involvement in thorax closure (Figure S4A) while rest eclosed as adults without thorax closure defect (Figure S4A).

A Mys-GFP reporter further revealed Mys enrichment at the HE-LEC layer contact points (Figure 4A-B) besides their enrichment at apico-lateral cell edges in both tissues (Figure 4B’, B” and their Y-Z optical sections). Further, knockdown of *mys* in both HE and LEC layer (see Figure S1B) or only the LEC layer (see Figure S1C) disrupted the process of thorax closure (double-headed arrow in left and middle panel in Video 6), culminating from the loss of LEC-HE contacts, besides the loss of LEC layer integrity, as seen from the loss of cell-cell contacts (yellow arrows in left and middle panel in Video 6, Figure 4C, D). By contrast, selective knockdown of *mys* only in the HE was marked by a characteristic snapping of the HE during the migration process from the LEC layer and culminating in a failed zipping of HEs (red arrow in the right panel of Video 6) besides detachment of the two cell layers (Figure 4E). These extreme phenotypic consequences of *mys* loss precluded adult emergence (Figure S4A). We thus compromised Gal4 expression by raising the larval cultures at a lower growth temperature (18°C) (see Material and Methods) (Duffy, 2002), which results in the eclosion of adults from ∼30% of these pupae, which invariably displayed cleft-thorax phenotype (Figure 4F-I). Additionally, live imaging from knockdown of *scb* in both LECs and HEs (Figure S4B) or LECs alone (Figure S4C), too, revealed an arrest of HE migration and loss of LEC layer integrity—like those seen following *mys* knockdown (Figure 4C-D). It is, therefore, likely that βPS1 (Mys) and αPS3 (Scb)—presumably as heterodimeric partners (Narasimha and Brown, 2004; Narasimha and Brown, 2013)—maintain LEC-HE adhesion—which is essential for the relay of pull forces from the LEC layer to HE.

**Figure 4:**
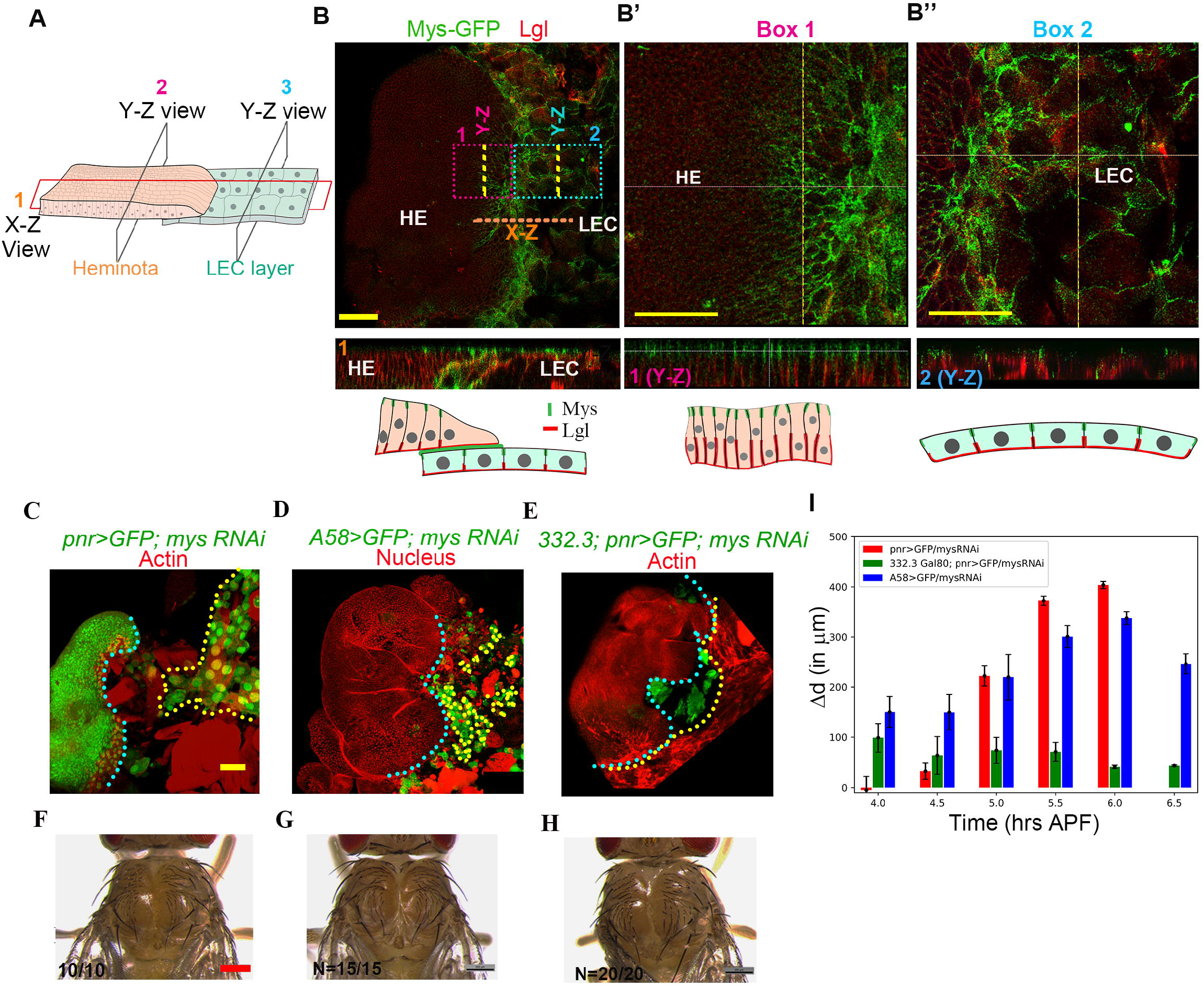
Integrin-βPS (Mys) is essential for maintaining HE-LEC contact as well as LEC integrity. **(A)** Cartoon representation to check Mys (Integrin βPS1) localization in both the tissues (2 and 3) and at the contact points (1, X-Z view) (Scale bar: 50 μm). **(B)**Mys*-*GFP localization (green) in the HE (magnified further in box 2), LEC (magnified further in box 3) and counterstained for Lgl (red). Broken yellow line marked as 1 to show the X-Z section for LEC-HE contact point and Y-Z view for box 2 and 3 were shown below for Mys localization with respective cartoon representation in (B) (Scale bar: A: 50 μm and Box 2 and 3: 25 μm). **(C-E)** Pupal whole mounts stained for actin (red) or nucleus (red): *mys*-knockdown using different Gal4s (see Figure S1B-D). The outlines for HE (cyan) and the LEC (yellow) are shown for their detachment (Scale bar: 50 μm) (also see Video 6). **(F-H)** Eclosed adults at lower temperature from C-E (18LC). Scale bar: 200 μm. **(I)** Deviation of HE separations at specific time points between control and *mys* knockdown (N = 4; Mean ± SEM, ANCOVA test was performed to compare the three linear regression models fits to the 3 cases; *** p values: <0.001).

### Epithelial dynamics in the LEC underlies thorax closure – an in-silico analysis

To test the biophysical underpinnings of this emergent mechanism of thorax closure, we utilized a two-dimensional vertex model of the LEC layer along with two flanking contralateral HEs. The vertex models have been utilized extensively to study epithelial mechanics in several developmental contexts, such as oriented cell divisions (Mao et al., 2011), density-independent phase transitions (Bi et al., 2015), epithelial topology (Aegerter-Wilmsen et al., 2010), cell flows during epithelial morphogenesis (Aigouy et al., 2010), cell size oscillations (Lin et al., 2017) or cell-cell adhesion mediated cell size dynamics (Kumar et al., 2020). In our vertex model, the mechanical response of the cells (LECs as well as HEs) is characterized by an energy function (Figure 5A) comprising of contributions from cell elasticity arising due to impermeability, or absence of leakiness of cell membrane, actomyosin-driven cell contractility and cadherin or integrin (at the interface of LEC layer and HE) mediated cell-cell adhesion (Farhadifar et al., 2007; Kumar et al., 2020). For overdamped cellular dynamics, the velocities of vertices are taken as proportional to the forces which are calculated from derivative of the energy functional relative to the vertex positions (Figure 5A, see Materials and methods).

**Figure 5:**
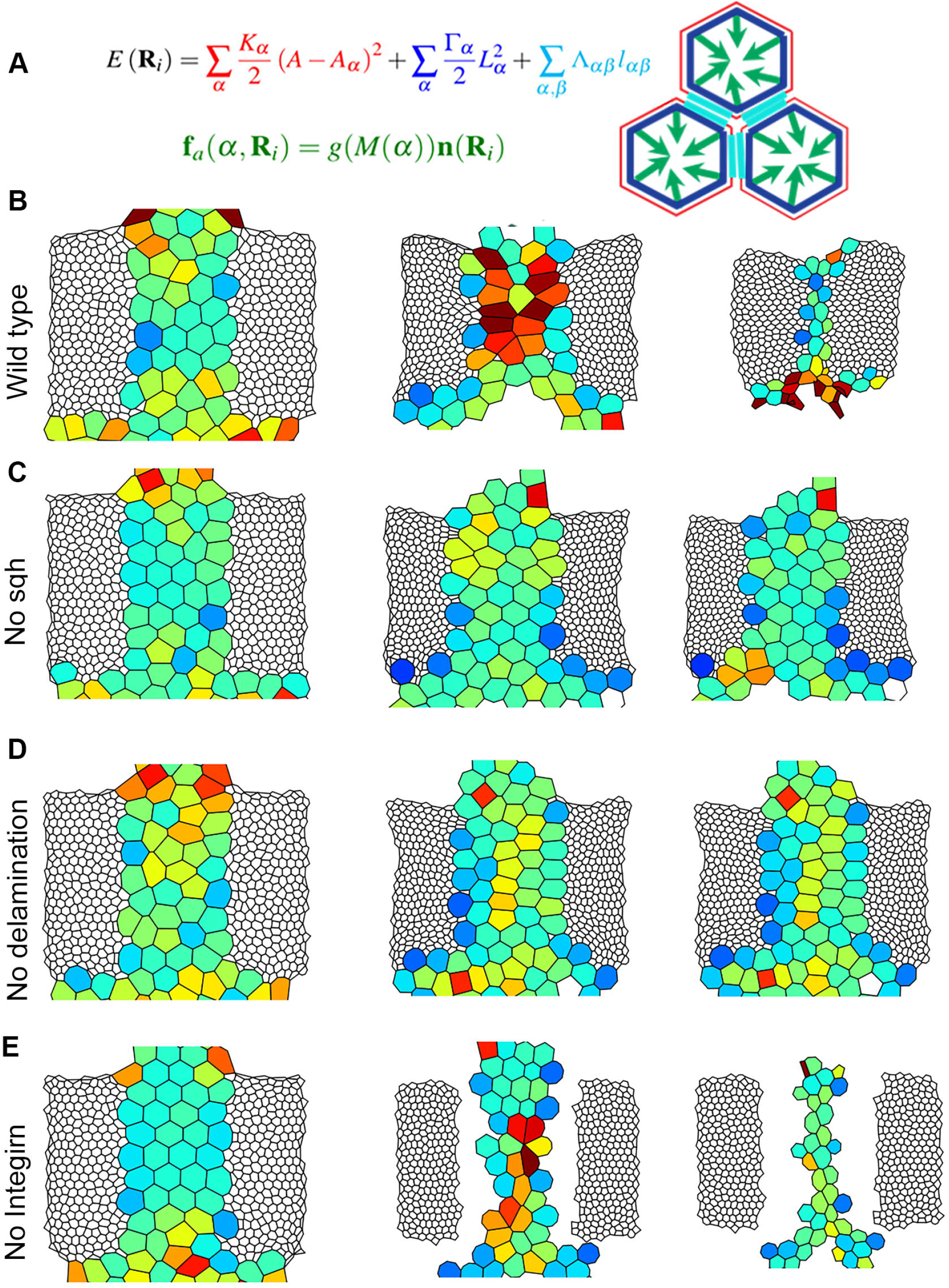
Mathematical modelling of LEC dynamics. **(A)** Energy functional describing the passive response of the epithelial tissue with contributions from cell elasticity (in red), actomyosin contraction (in blue), and cell-cell adhesion (in cyan). Sqh localization at the cell membrane is considered in the form of active forces (in green). **(B-D)** Three snapshots showing the cell geometries in the initial, intermediate and late stages of thorax closure in wild type scenario. Same snapshots for (C) in the absence of active force, analogous to *sqh-RNAi*, and (D) when the cell delamination is blocked, and (E) when integrin is knockdown.

**Figure 6:**
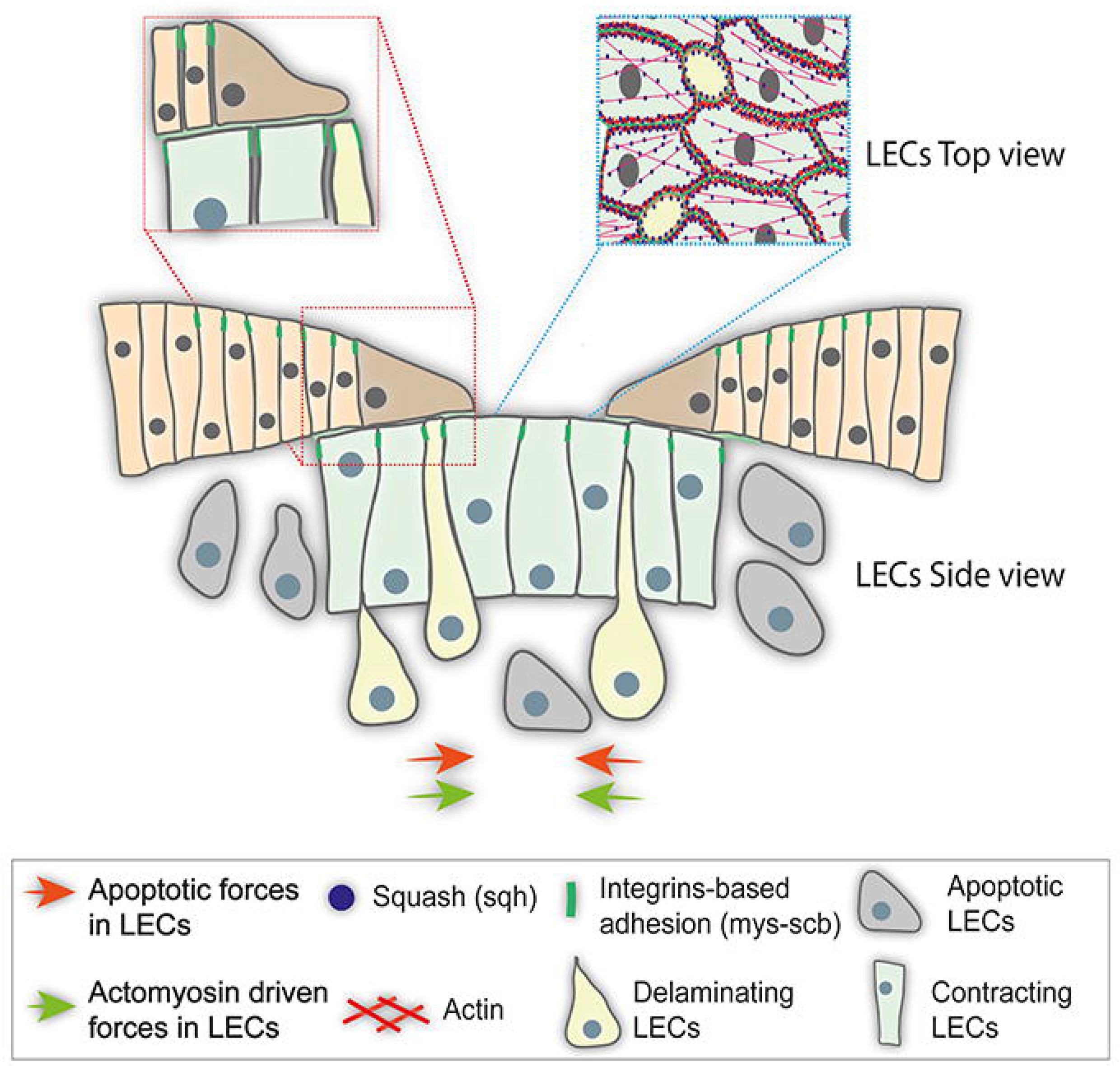
LEC layer generate pull force during thorax closure. Graphical summary of the LEC layer (green) dynamic during thorax closure. LEC layer displays two different force generating mechanisms: a) via virtue of their enriched actomyosin network that allow cells to undergo active contraction (green arrow), and b) crowding induced delamination (rosette yellow cells) (orange arrow), together providing a pull force that drags the above HE to come close at the body midline. Further, LEC layer and HE edge cells are indirectly connected via integrin-based connections (Mys-Scb heterodimer, green), besides LECs displaying integrin-enrichment at the sub-apical junctions, that allow relay of the LECs-forces to the HE tissues and stability to the LEC layer during the contraction phase.

The numerical simulation of the vertex model of LEC and HE recapitulates the experimental observations, as shown (first panel in Video 7, Figure 5B). Starting from irregular cell geometries in the LEC layer, cells are seen to elongate along the L→M direction due to higher drag offered by the HE (Figure 5B), reminiscent of such transitions seen *in vivo*. Further, in the absence of active forces, mediated by Sqh and delamination, in the LECs *in vivo* resulted in arrested thorax closure (second and third panel in Video 7, Figure 5C and 5D). Finally, lowering of cell contacts between LECs and HEs—representing the fallout of *in vivo* knockdown of integrins in HE-specific manner (see Figure S1C) (fourth panel in Video 7)—arrested HE migration and thorax closure (Figure 5E). These findings from vertex simulation, therefore, further reaffirm that LEC delamination and contraction-mediated pull forces relayed to HE are essential biophysical principles of thorax closure.

## Discussion

A comparison between embryonic DC and pupal thorax closure reveals uncanny parallels between their respective LEs: both leading their respective epithelia all the way to migrate over a degenerating cell layer of a preceding developmental stage (Belacortu and Paricio, 2011; Zeitlinger and Bohmann, 1999). Given the primary importance assigned earlier to the LEs for the migration of embryonic lateral epidermis during DC (Kiehart et al., 2000), it may appear reasonable to assume LEs as a force-generator during thorax closure (Martín-Blanco et al., 2000). Instead, our results reveal that it is the LEC layer generated pull forces that are necessary during the thorax closure. During thorax closure, the LEC layer displays elaborate orchestration of epithelial dynamics marked by squamous-to-pseudostratified columnar transition and accompanying the non-muscle myosin-II-mediated contractions, cell extrusion and cell death. Pull forces, thus generated in the shrinking LEC layer, are then relayed to the flanking HEs via their mutual integrin-based adhesions (Figure 6). From the biomechanical viewpoint, too, our simplified two-dimensional vertex model formalizes these essential elements of the LEC layer in thorax closure: namely, their cell delamination, non-muscle myosin-II driven contractions and finally its integrin-based adhesion with HEs (Figure 5). In essence, therefore, reminiscent of embryonic amnioserosa ((Ducuing and Vincent, 2016; Pasakarnis et al., 2016) for review see, (Hayes and Solon, 2017; Kiehart et al., 2017)), the degenerating thoracic LEC layer, too, generates the forces required to drive its replacement by successors—the HEs.

### LEC generates the pull forces for HE migration

Our results reveal two core attributes of the LEC layer, which underlie HE migration. The first attribute is its non-muscle myosin II-mediated contractions. It is conceivable that actomyosin-mediated contraction of the LECs, too, contribute to their cell shape remodeling, reminiscent of other morphogenetic processes, such as gastrulation, oogenesis, and intercalation during germband elongation (for review, see (Heisenberg and Bellaïche, 2013)). For instance, deep tissue imaging of *Drosophila* gastrulation reveals that apical actomyosin meshwork drives apical contraction in the ventral furrow cells leading to cell lengthening and basal movement of the nuclei: the net fallout being volume conservation within a constrained space (Gelbart et al., 2012; Khan et al., 2014). During thorax closure, too, the LECs display apical contractions (see Video 2, Figure 2A) and cell elongation (Figure 1I and K-L)—accompanying their squamous-to-columnar epithelial transition—which could also mean their volume conservation during cell shape remodeling (for review (Gorfinkiel, 2016)). Further, the apical enrichment of actin and Sqh (Figure 3D-D’’’) in the LEC layer during thorax closure not only generates pull forces but could as well be responsible for an increase in cell density. A consequent cell overcrowding in the LEC layer on the other hand is offset by cell extrusion and cell death (Figure 6). This essential attribute of the LECs are also seen in amnioserosa during DC, marked by cell extrusions (ingression) via apoptosis ((Toyama et al., 2008) for review, see (Ambrosini et al., 2017; Kiehart et al., 2017; Teng and Toyama, 2011)). In other migrating epithelia, too, such as the HEs, cell overcrowding-mediated cell extrusion alleviate space constraint, leading to restoration of cell number homeostasis ((Levayer et al., 2016; Marinari et al., 2012) for review, see (Ambrosini et al., 2017; Fadul and Rosenblatt, 2018)).

The second core attribute of the LEC layer is its integrin-based adhesion with the flanking HEs that allow relay of the pull forces to the latter (Figure 6). This scenario, too, appears to recur in multiple contexts of animal development, including DC in *Drosophila* embryogenesis (Muliyil et al., 2011; Narasimha and Brown, 2004) or in mouse embryos wherein surface ectoderm connecting contralateral neuroepithelium—the precursor to the neural tube—display enriched levels of integrin β1 subunit apically (Molè et al., 2020).

### Thorax closure: an archetypal mechanism for tissue or wound closure

One of the enduring revelations of various tissue closure mechanisms during development or tissue repair is the critical role of the cell death in the degenerating cell layer. For instance, during wound healing, injured and dying cells send out a signal—both humoral (cytokine) and mechano-stimuli—to the surrounding healthy cells to migrate and proliferate and thereby leading to their replacement (for review, see (Eming et al., 2007; Gurtner et al., 2008; Martin, 1997; Shaw and Martin, 2009)). For instance, a wounded site is marked by aggregation of myofibroblasts, which represent transformed fibroblasts (Alberts et al., 2002) or are derived from epithelial cells through EMT (Yang and Liu, 2001). These are enriched in stress fibers and thus generate contractile forces to bring the wound edges closer. Upon wound healing, myofibroblasts are eliminated by apoptosis (Darby et al., 2014; Li and Wang, 2011). LECs, too, display functional parallels with myofibroblasts at wound sites. For instance, while these LECs remain dormant during larval life, their secretory activities are heightened upon pupariation and secretes of pre-pupal cuticle (Fristrom and Fristrom, 1993). Further, reminiscent of fibroblasts-to-myofibroblasts transition upon tissue injury, abdominal LECs, too, display EMT-like transition marked by filopodial and lamellipodial protrusions (Arata et al., 2017; Bischoff, 2012). These uncanny parallels between LECs *in situ* during thorax closure and the myofibroblasts, which migrate at the site of tissue closure, further show that the principles of thorax closure revealed here are primitive, presumably predating the emergence of holometabolous insects, and have been co-opted for multiple contexts of developmentally regulated tissue closure and repair events.

## Materials and Methods

### Fly stocks and rearing conditions

Stocks used in this study include:

#### Gal4 lines

*pnr-Gal4* (Chr. III) (Calleja et al., 2000) and *A58-Gal4* (Chr. III) (Galko and Krasnow, 2004).

#### Gal80 lines

*332*.*3Gal80* (Chr. II) (Pasakarnis et al., 2016).

#### UAS-based reporter lines

*UAS-(nls)GFP* (Chr. III) (BDSC #4776), *UAS-nuDsRed* (Chr. III) (Galko and Krasnow, 2004), and *UAS-Apoliner* (Chr. II) (BDSC #321122) (Bardet et al., 2008).

#### UAS-overexpression lines

*UAS-p35* (Chr. X) (BDSC #6298).

#### UAS-knockdown lines

*UAS-sqh-RNAi* (Chr. III) (BDSC #33892), *UAS-Itgbn-RNAi (*βυ) (Chr. III) (BDSC #28601), *UAS-mys-RNAi (*β*PS)* (Chr. III) (BDSC #27735), *UAS-mew-RNAi (*U*PS1)*(Chr. III) (BDSC #27543), *UAS-if-RNAi (*U*PS2)* (Chr. III) (BDSC #27544), *UAS-scb-RNAi (*α*PS3)* (Chr. III) (BDSC #27545), and *UAS-ItgaPS4-RNAi (*α*PS4)* (Chr. III) (BDSC #28535).

#### Reporter lines

*puc-LacZ*^*E69*^ (Chr. III) (Martín-Blanco et al., 1998), *mys-GFP* (Chr. X) (Klapholz et al., 2015), *sqh-GFP* (Chr. II) (BDSC # 57145), *lac-GFP* (BDSC #6833) (Morin et al., 2001) and *spider-GFP* (Chr. III) (BDSC #59025) (Morin et al., 2001).

All fly cultures, unless mentioned otherwise, were grown at 25±1°C on standard fly food. Knockdown or overexpression-based studies were performed on 25±1°C. For *mys* adult studies, second-instar larvae were transferred to 18°C to decrease Gal4 activity (Duffy, 2002) and rescue high pupal lethality.

### Staging, dissection, and immunostaining of pupal HE

0 hrs APF pupae were carefully selected and dissected at desired time points, as described previously (Kumar et al., 2020). In brief, dissection was performed and fixed in 4% PF in 1X PBST followed by 1X PBST washes and antibody staining. Primary antibody and their dilutions used were: anti-Lgl (1:50) (Gift from Fumio Matsazaki) and β-galactosidase (1:200) (raised in mouse; Sigma Aldrich, Catalog No: G8021). Secondary antibodies used were tagged with Alexa 488, 555, or 633 (Molecular Probes). Phalloidin (488, 555 or 633) and ToPro (555 or 633) were used to mark F-actin filaments and nucleus, respectively. Final mounting was done in Vectashield (H-1000, Molecular Probe) using a bridge slide.

### Pupal cryo-cross section preparations

For the pupal cryosections, staged samples were fixed overnight with 4% PF in 1X PBS at 4°C followed by washing in 0.1% PBST and treatment with graded sucrose (10, 20, and finally in 30%). Later, the samples were treated in an OCT (Leica Biosystems, Product No. 3801480) and 30% sucrose media (1:1) before final mounting in the OCT followed by freezing. 15 μm thick sections were cut using Leica cryotome, counterstained, and mounted in lysine coated glass slides.

### Image acquisition and processing

For whole-mount and cryosection preparations, confocal images were acquired using the Leica SP5 laser scanning confocal microscope. Pupal (still and time-lapse) and adult imaging were performed using Leica M205FA or Zeiss AxioZoom V1.6 microscope on either bright field or GFP filters. Images were processed using LAS 4.0, ImageJ software, and Adobe Photoshop CS6, whichever relevant.

### Laser ablation in LEC layer

A modified FRAP protocol was used in a tunable infrared laser attached to Leica multiphoton SP8 confocal microscope to ablate the LEC layer. Initially, the staged pupae (4:30 hrs APF) was imaged for 30 mins (30 seconds per frame) to reach 5 hrs APF and shifted to FRAP conditions where 100% laser at 900 nm was focused in a narrow rectangular ROI (red box, Figure 3B) on the LEC layer for 60 secs and imaged again in normal conditions for 15 mins (30 sec per frame) (Video 3).

### Calculation of inter-HE distance

Inter-HE distances during various perturbations was calculated based on measuring the L→L distance between migrating leading edges of the HE. The graph was then plotted in GraphPad 8.0.

### Quantification of various LEC layer parameters

#### LEC surface area

For the measurement of the LEC layer surface area over time, outlines were manually drawn along the perimeter of the LEC layer with Nu-DsRed marked nucleus (*A58>nu-DsRed*) on selected time-lapse images during thorax closure. The surface areas were then measured using ImageJ software.

#### LEC nucleus number

For counting the nuclear number in the LEC layer, live imaging video of Nu-DsRed marked nuclei (*A58>Nu-dsRed*) (right panel in Video 1) was used, and nuclei numbers were counted using the particle analyzer tool in ImageJ.

#### LEC layer crowding

Three boxes of equal size were placed in the medial LEC layer (Box 1 and 2 on T2 and Box 3 in T3 segments), followed by manual counting and plotting in Excel.

#### LEC nuclear migration

Each frame of the right panel in Video 1 with *A58>Nu-dsRed* stained nuclei was smoothened using a Gaussian filter followed by image thresholding. The location of each nucleus in each frame was subsequently detected by finding the local maxima in the image with skimage Python library. For movie frames corresponding to 0-4 hrs APF and 4-6 hrs APF, the location of the nucleus was plotted using scatter plots, in blue and green colors, respectively.

#### LEC layer thickness

Y-Z sections were used to calculate the thickness of the LEC layer (*pnr>GFP)* of pupae at two different time points, before and during thorax closure. Image analysis was performed on LAS-X software followed by their plotting on GraphPad Prism 8.0.

#### Mathematical model

##### Introduction to vertex model of epithelium

Vertex modeling is a handy tool for the study of the biophysical mechanisms in the epithelial tissues, which is a three-dimensional structure, by the representation of apical cross-sections of the cells with an assumption that the cell geometry does not change as one moves from the apical to the basal end of a cell. Vertex modeling permits the testing of the essential elements of the LEC dynamics and its relay to the HE in a highly simplified version posed side by side (Bi et al., 2015; Farhadifar et al., 2007; Kumar et al., 2020). Further, the epithelial dynamics has been considered in the vertex model to be intrinsically regulated based on the features embedded in itself, and thus do not factor the precise trajectories of LEC shrinkage displayed by LEC apolysis and during the thorax closure proper (see Figure 1). Further, confluent LEC and heminota in a monolayer in our vertex model is not an exact representation of these tissue alignments seen *in vivo*, where these are seen in different planes. Furthermore, the two-dimensional vertex-based representation also does not consider the contributions other than that of LEC and HE cells, such as thorax geometry, the effect of the larval segment boundaries, or even the activity of the muscular layer underlying the LEC layer. These simplifications notwithstanding, vertex modeling allows substantial resolution of the biophysical underpinning of the LEC dynamic and its fallout on the HEs.

In this reduced two-dimensional framework, the mechanics of an epithelial monolayer is described by the energy functional:

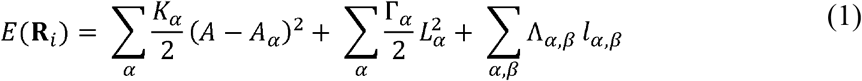

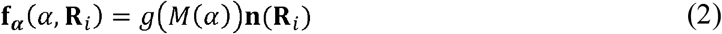

where *K*_α_ represents the modulus of area elasticity for cell-α, which is, in this context, a combined manifestation of the incompressibility of cytoplasm and impermeability of the cell membrane, Λ_α β_ incorporates the cadherin dependent cell-cell adhesion at the edge shared between two adjacent cells α and β, and, finally, Г_a_ stands for cell contractility arising due to the actomyosin network near the apical end of the cell. Following the approach proposed earlier, we can non-dimensionalize the adhesion parameter and contractility by 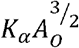 and *K*_α_*A*_o_ respectively (Farhadifar et al., 2007).

From the energy functional, one can calculate the force acting on each vertex by taking the functional derivative of the above equation for the vertex coordinates. Under the assumption of overdamped dynamics, the velocity of each vertex is proportional to the total force acting on it.

###### T1 transition

In addition to the vertex movements due to the forces acting on it, in this model T1 transition is also taken into account where a collective of four cells exchange some of their neighbors (see Figure M1A). The T1 transition occurs whenever the edge, shared between any two cells, becomes smaller than a pre-specified threshold value.

###### T2 transition

Further, whenever the two-dimensional apical cross-section of a cell becomes smaller than a pre-specified threshold value, the cell is considered to get detached from the epithelial monolayer (see Figure M1A). It can be understood by the fact that a minimal cell in the vertex model, which is a two-dimensional representation of apical end only, implies that most of its contents are already out of the plane of the tissue, and the cell is destined to get delaminated from the tissue. This process is known as the T2 transition.

**Figure M1:**
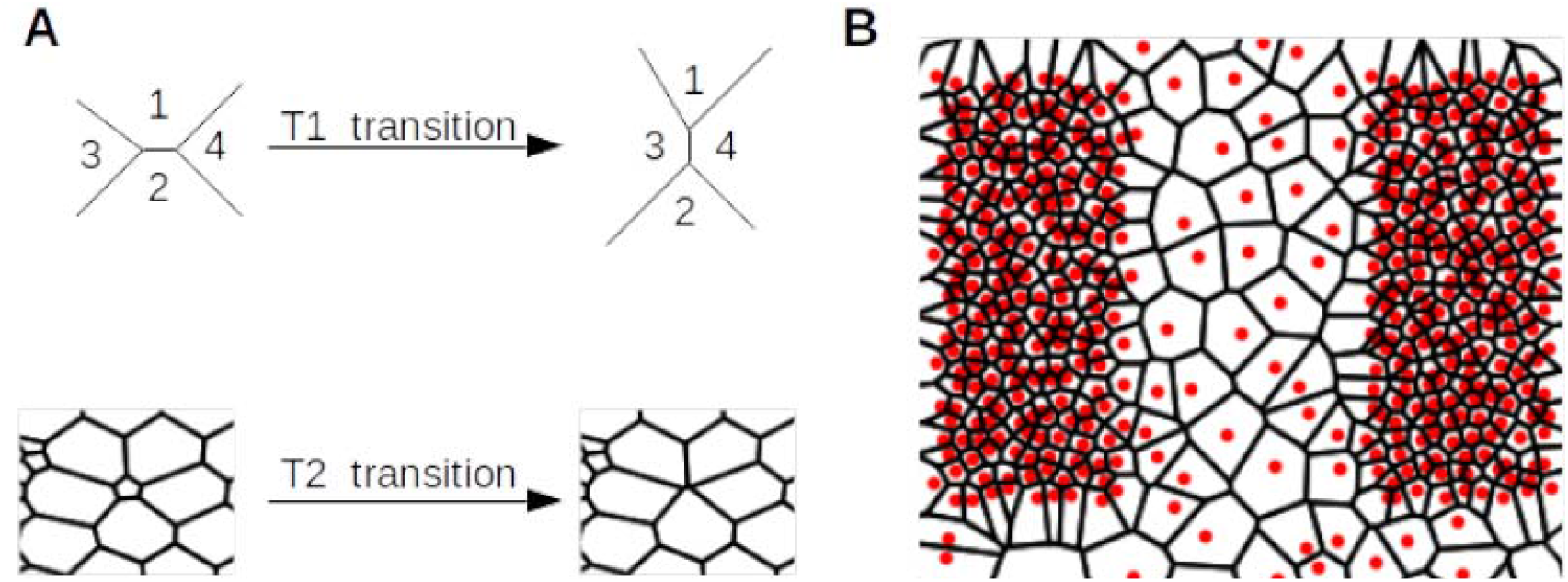
**(A)** Schematic depicting T1 and T2 transitions. **(B)** The generation of initial cell configurations for the LEC layer and HE. The red dots denote the initial seeds for the Voronoi tessellation with sparse and dense red dots representing LEC and HE, respectively.

With this description of the basic vertex model (without any tissue-specific ingredients) in the following, we will describe the tissue-specific model considerations.

##### Initial cell geometry generation

For the study of the LEC dynamics, we utilize the same framework while considering both of the LEC and two flanking contralateral HEs. It is worth recalling here that even though both the tissues are epithelial, their detailed descriptions differ significantly since LECs are of squamous characteristics (large apical section), and HE cells are of columnar type (smaller apical section). To generate a monolayer of LEC with HE for the vertex model, we use Voronoi tessellation with the seed points with higher and lower densities in the HE and LEC regions, respectively (see Figure M1B).

##### Vertex model for HE

For the modeling of the HE cells, we utilize the vertex model without any additional consideration. Although, as reported in one of our earlier works, the dynamics of the HE cells is regulated by the expression gradients of members of the planar cell polarity pathway, we do not take that detail into account here (Kumar et al., 2020). We do not expect the findings to change even if those details are incorporated into the model.

##### Vertex model for LEC layer

For the LEC layer, we extend the vertex model to incorporate the role of the Sqh, as observed in the experiments. We take into account the contractile activity of the Sqh in the form of active forces on the boundaries of the LEC cells in the direction normal to the cell membrane. The active force on the *i*^*th*^ vertex of the cell is:

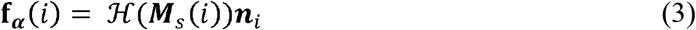

where, ℋ is the Hill’s function, ***M***_*s*_ is the level of Sqh in the cell and is ***n***_*i*_ the membrane inward normal at the *i*^*th*^ vertex. This nature of active forces has previously been used to model motile cells of amoeboid nature (Farutin et al., 2013). It has to be noted that due to the intrinsic nature of the active force, the total active forces and total torque due to the active force on the cell should vanish. That is:

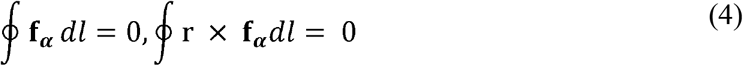

where, the integrals are along the cell perimeter. In order to ensure this, we add a correction to the active forces in the form:

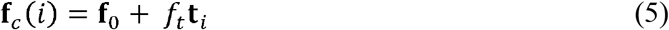

where t_*i*_ is the unit vector tangent to the cell membrane. The exact values of the and are calculated from equation (4). These corrections ensure that the total active force and torque on each cell vanish (Wu et al., 2016). For the LECs, we considered that the level of Sqh is equal in all the cells. To keep the computer simulation of the model, we apply active forces in a ramping manner starting from 0 to a fixed maximum value in linear increment over time.

### Boundary conditions

Forces from the boundaries play a significant role in the overall dynamics of the epithelial tissues (Cochet-Escartin et al., 2014; Nikolić et al., 2006) and are so expected in thorax closure. In the simulation of thorax closure, we have considered all the boundaries to be stress-free. It needs to be pointed out that this boundary condition represents a simplification of the *in vivo* condition where the tissues and geometric features, such as the segment boundary, adjacent to the LEC or HE may present additional constraints to the cell dynamics.

### Simulation of different cases shown in Figure 5

Figure 5 shows the results of vertex modeling of the thorax closure with the following cases:

1. The Wild-type case (Figure 5B) considers all the aforementioned model ingredients.
2. For the simulation of LEC layer dynamics with no Sqh (Figure 5C), we reduced the value of Sqh to 1/10^th^ of the Wild type case.
3. For the simulation with no cell delamination (Figure 5D), the T2 transitions were not performed.
4. For the simulation with integrin loss (Figure 5E) the adhesion between the LECs and the HE cells was broken.

## Acknowledgment

We would like to thank Damein Brunner for *332*.*3Gal80*, Michael Galko for *A58-Gal4* and *UAS-nudsRed*, and Nick Brown for *mys-GFP* stocks, and Fumio Matsuzaki for the antibody (Lgl). We would also like to thank Richa Rikhy for her guidance in conducting Laser ablation experiments in IISER Pune, and Maithreyi Narasimha, TIFR Mumbai and Nick Brown, University of Cambridge for their inputs in our work. We further thank Suvimal Kumar Sindhu and Jonaki Sen for their help in tissue cryo-sectioning.

## Supplementary Figure Legends

**Supplementary Figure 1:**
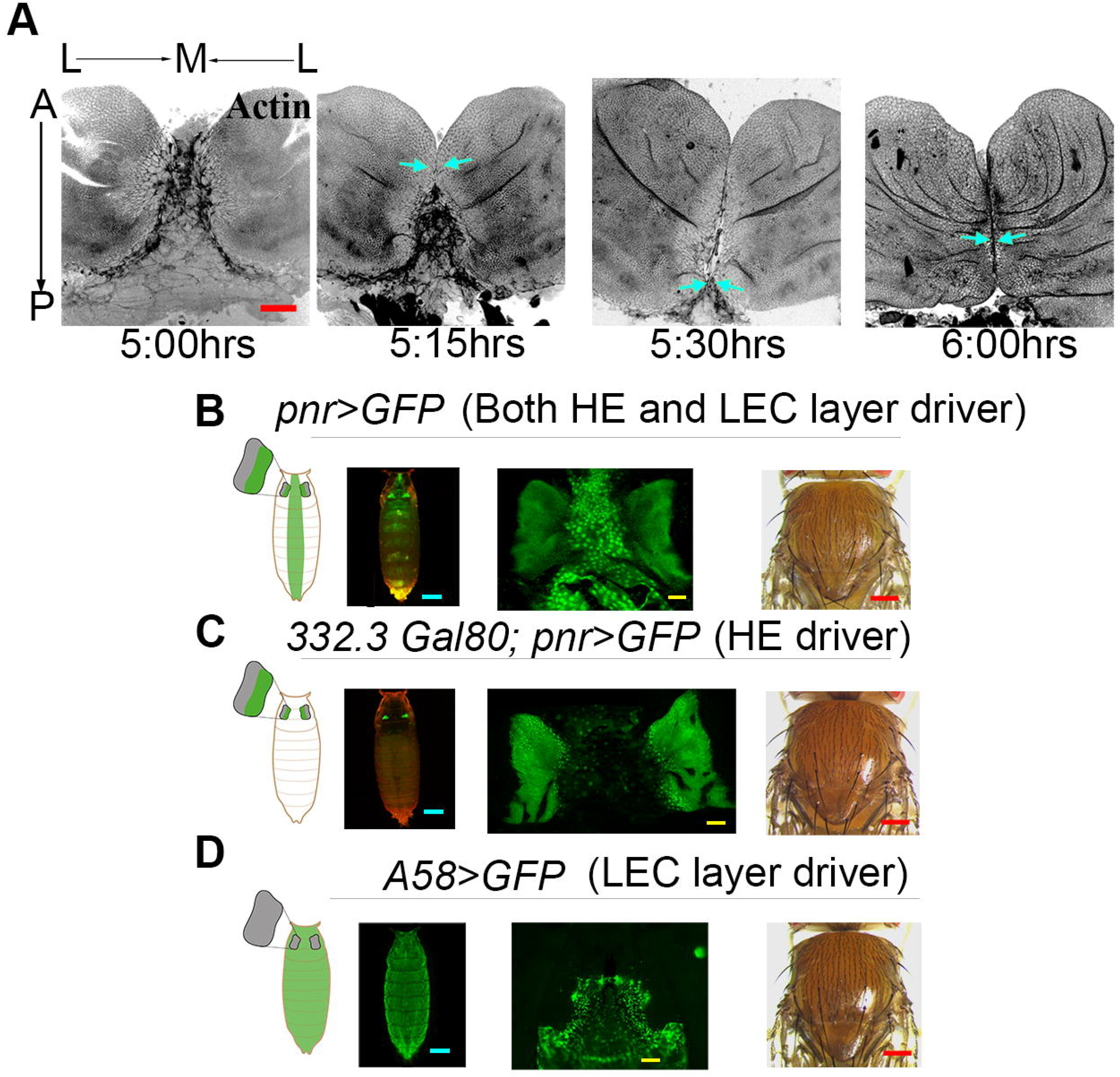
HE zippering event and the tissue-specific Gal4s used in current study. **(A)** Contralateral HEs, undergoing fusion (blue arrow), with LEC layer, marked for actin (grey) at late stages of thorax closure (i.e. 5:00, 5:15, 5:30, and 6:00 hrs APF). Scale Bar: 50 μm. **(B-D)** Domains of expression of *nls-*GFP driven by *pnr-Gal4* (B); *332*.*3Gal80; pnr-Gal4* (C) and *A58-Gal4* (D). *332*.*3Gal80* suppresses the expression of *pnr* in LECs. Cartoon represent (column 1), actual view in pupae (column 2) and in dissected HE and LEC layer (column 3) with their eclosed adults (column 4) (Scale bars: pupae: 500 µm, tissue mounts: 50 µm and adults: 200 µm).

**Supplementary Figure 2:**
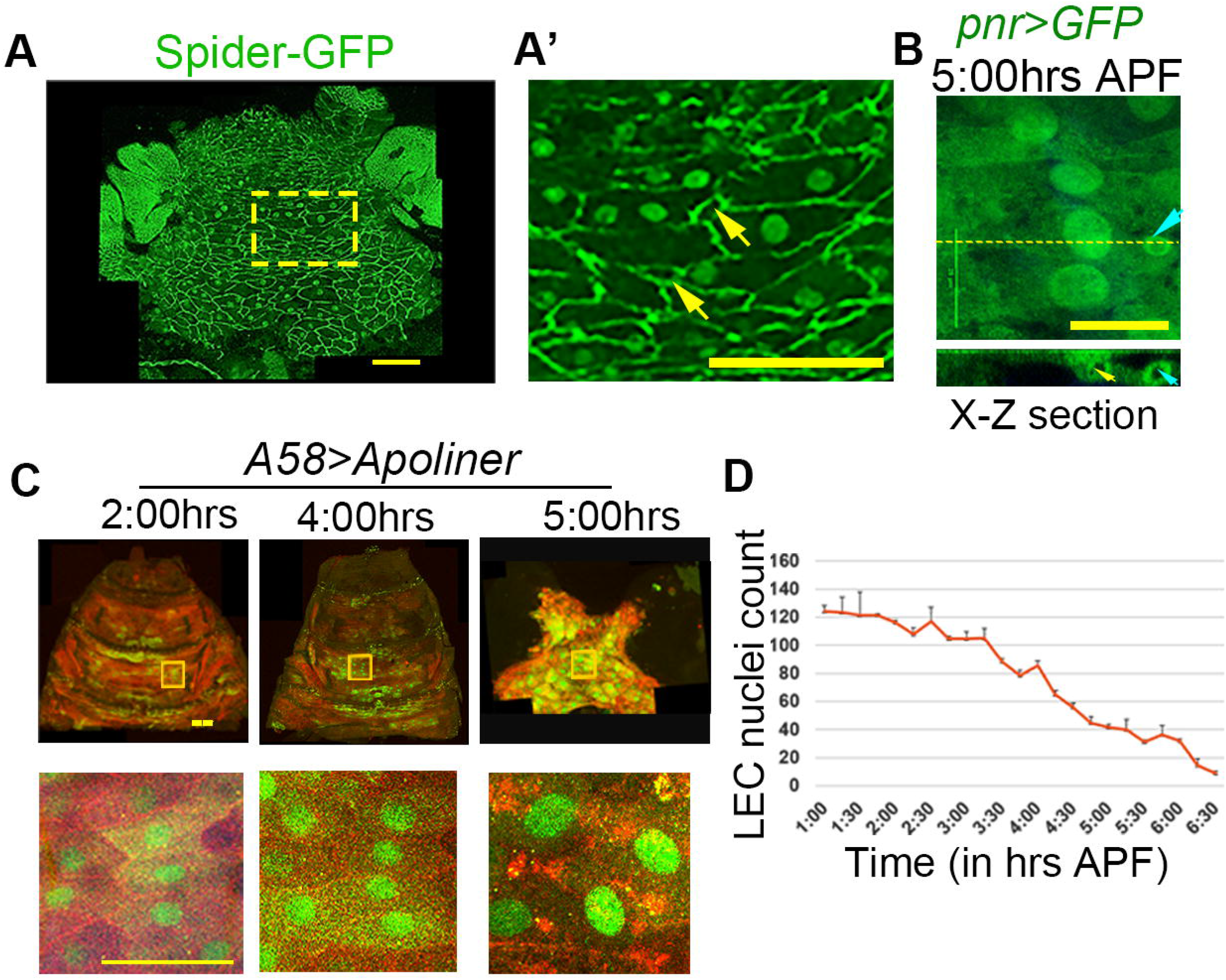
LECs shows delamination and early caspase activation. **(A-A’)** Whole mount preparation of Spider-GFP marked LECs that shows the presence of delaminating cells—marked by their small apical sizes. **(B)** X-Z view of LECs *(pnr>GFP)* to show delaminated cells (blue arrow) at 5:30 hrs APF. Scale bar: 25 μm. **(C)** Apoliner expression (green nuclei) before (2 hrs APF) and during (4 and 5 hrs APF) thorax closure. Boxed area from each image are shown at a higher magnification in the lower panel (Scale bar: upper panel: 100 μm, lower panel: 50 μm). **(D)** Loss of cells—counted using number of nucleus—in LEC layer (from T2 segment) during the thorax closure (N=2, Mean ± SEM).

**Supplementary Figure 3:**
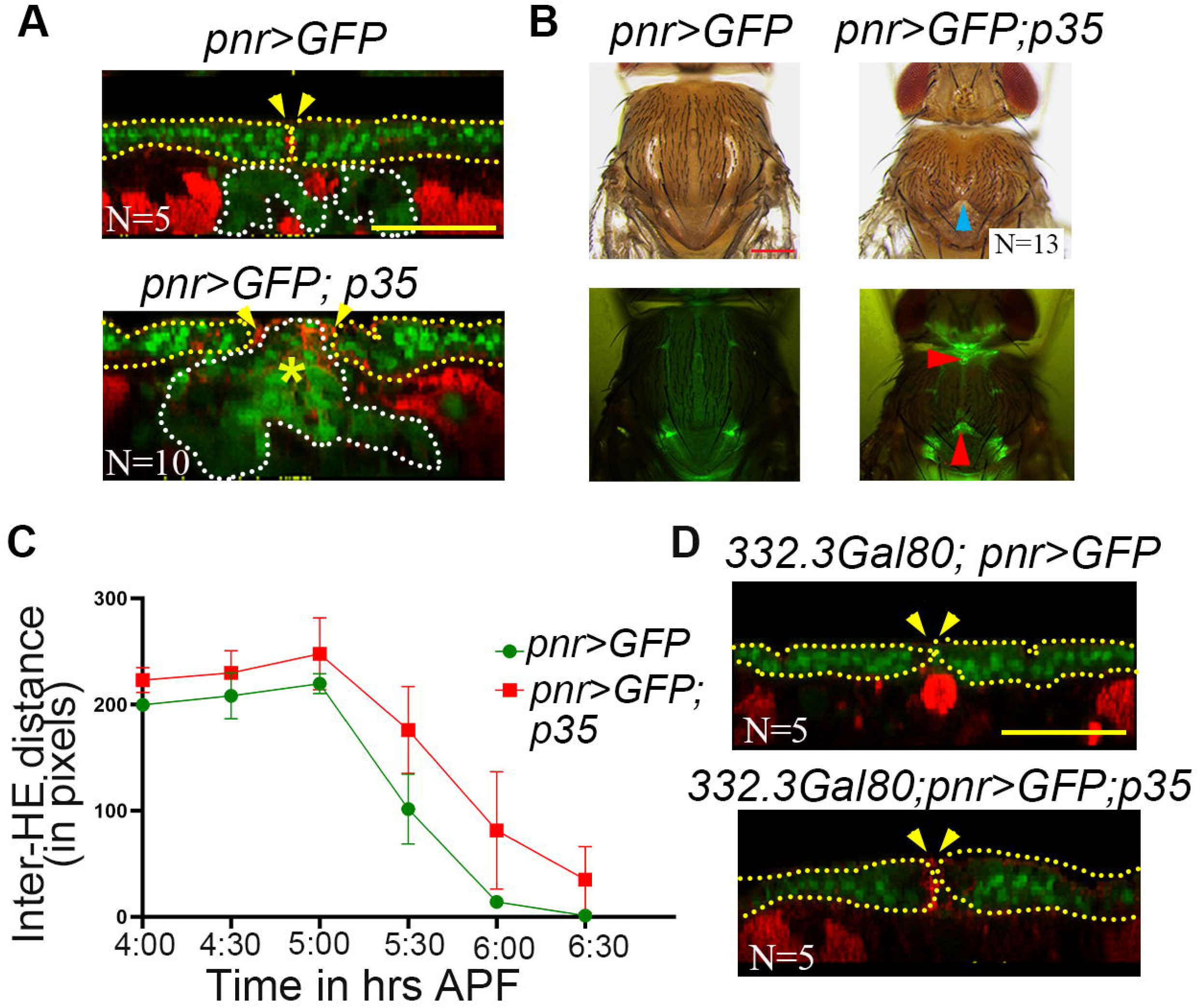
p35 overexpression spanning LECs impediment the process of HE migration and fusion. **A)** Comparison of the X-Z view for the *pnr-Gal4* driven p35 overexpression (lower panel) with respect to its control (upper panel), counter stained for actin (red). **(B)** Adult thorax (upper panels) and their respective fluorescent images (lower panels) from the above cases. Cleft thoraces phenotypes following *p35* overexpression (blue arrowheads in upper panel) and red arrowheads mark persistence of GFP-marked LECs in adults (lower panel) (Scale bar: 200 μm). **(C)** Inter-HE distances in case of *pnr-Gal4* and their respective *p35* overexpression (N=3 for each case, Mean ± SEM, two-way ANOVA analysis was performed with **** p value:<0.0001). **(D)** Comparison of the X-Z view for the *332*.*3Gal80; pnr-Gal4* driven p35 overexpression (lower panel) with respect to its control (upper panel), counter stained for actin (red).

**Supplementary Figure 4:**
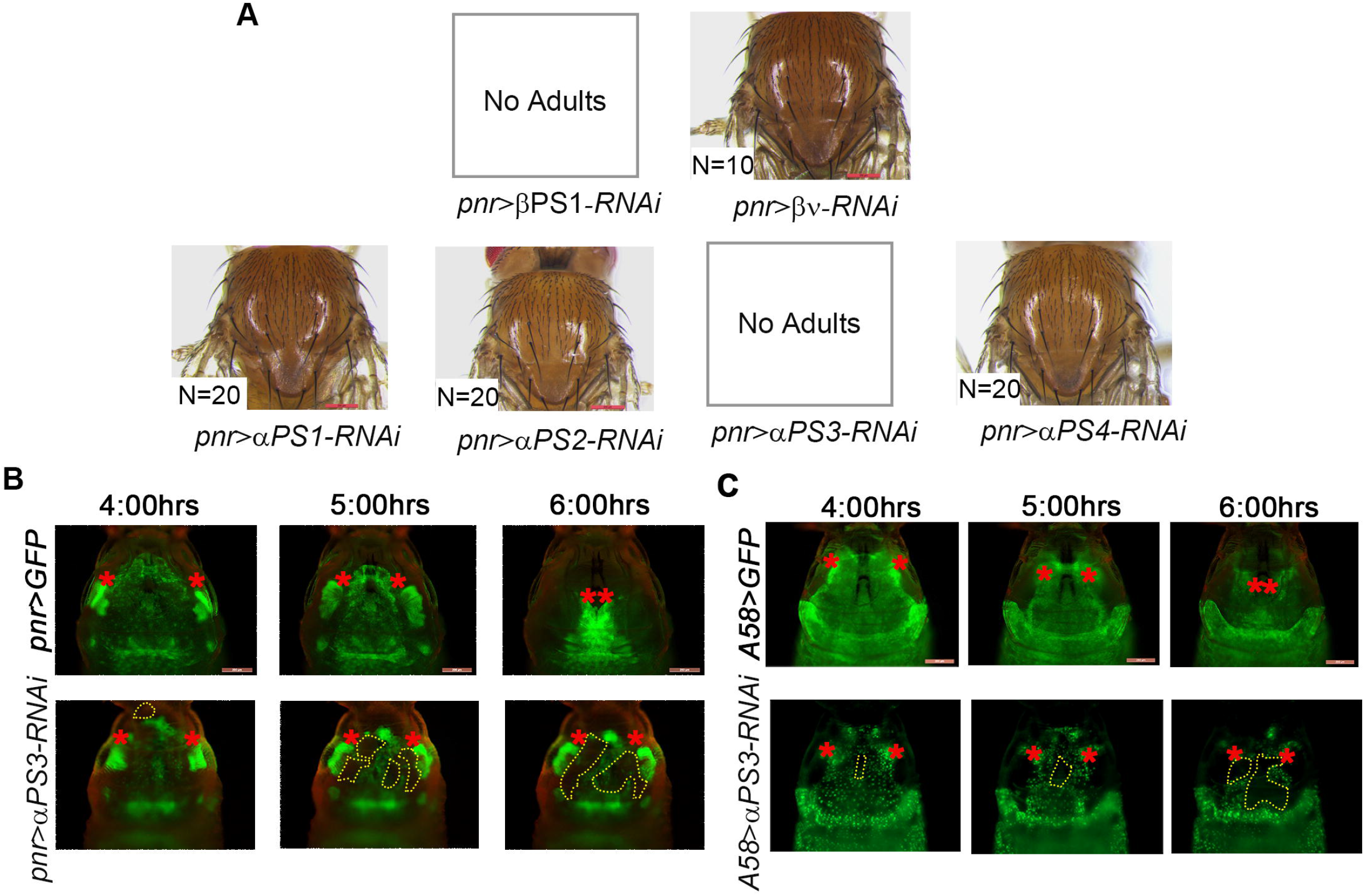
*scab*, □-PS3 integrin, cooperates with *mys* during the process of thorax closure. **(A)** Screening for the candidate integrin heterodimer involved during thorax closure. In case of knockdown of βPS1 (*mys)* and αPS3 (*scb)* results in no adults whereas in rest of cases eclosed adults are without any thorax defect. Scale bar: 200µm. **(B-C)** Time-lapse snapshots at different time intervals of thorax closure where αPS3 (*scab*) is knocked down using *pnr-Gal4* (E) and *A58-Gal4* (F) (lower panels) as compared to their respective controls (upper panels). Heminotal position in each case is marked in red star.

## Video Legends: (see supplementary information)

**Video 1_Live imaging of LECs and HE:** Left panel showing dual tissue live-imaging during early, mid and late stages of HE migration and finally fusion, whereas, right panel shows the LECs nuclear migration tracks before and during thorax closure.

**Video 2_Apical cell constriction and delamination in LEC layer:** Lachesin-GFP marked LECs boundaries showing apical constriction (yellow) and delamination (red).

**Video 3_Laser ablation of LEC layer:** *pnr>GFP* staged pupae showing events before and after the laser ablation (red arrow).

**Video 4_Domains of Gal4 driver expression**: Controls used for live imaging to understand the process of thorax closure, *pnr>GFP* (left panel), *A58-GFP* (middle panel) and *332*.*3Gal80; pnr>GFP* (right panel) are used in this study.

**Video 5_Tissue-specific *sqh* knockdown:** Live imaging of pupae when *sqh* is knockdown using different Gal4 mentioned above: that is, *pnr>GFP; sqh-RNAi* (left panel), *A58-GFP; sqh-RNAi* (middle panel) and *332*.*3Gal80; pnr>GFP; sqh-RNAi* (right panel).

**Video 6_Tissue specific *mys* knockdown:** Live imaging of pupae when *mys* is knockdown using different Gal4 mentioned above: that is, *pnr>GFP; mys-RNAi* (left panel), *A58-GFP; mys-RNAi* (middle panel) and *332*.*3Gal80; pnr>GFP; mys-RNAi* (right panel).

**Video 7_Mathematical simulation of various LEC parameters during thorax closure:** Visualization of thorax closure when all LEC parameters, identified in this study, are considered (Wild-type, first panel), when non-muscle myosin-II component is absent (second panel), when delamination is absent (third panel), and when integrin adhesions between the two-tissue interface are absent (fourth panel).

